# Adaptive evolution of olfactory degeneration in recently flightless insects

**DOI:** 10.1101/2020.04.10.035311

**Authors:** Stefanie Neupert, Graham A. McCulloch, Brodie J. Foster, Jonathan M. Waters, Paul Szyszka

## Abstract

Fast-moving animals need fast-acting sensory systems. Flying insects have thus evolved exceptionally quick visual (1) and olfactory processing ability (2). For example, flighted insects can track the temporal structure of turbulent odor plumes at rates above 100 Hz (3). The evolutionary lability of such sensory systems, however, remains unknown. We test for rapid evolutionary shifts in olfactory processing speed associated with flight loss, through neurobiological comparisons of sympatric flighted versus flightless lineages within a wing-polymorphic stonefly species. Our analyses of sensory responses reveal that recently-evolved flightless lineages have substantially degraded olfactory acuity. By comparing flighted versus flightless ecotypes with similar genetic backgrounds (4), we eliminate other confounding factors that might have affected the evolution of their olfactory reception mechanisms. Our detection of different patterns of degraded olfactory sensitivity and speed in independently wing-reduced lineages highlights parallel evolution of sensory degeneration. These reductions in sensory ability also echo the rapid vestigialization of wings themselves (4, 5), and represent a neurobiological parallel to the convergent phenotypic shifts seen under sharp selective gradients in other systems (e.g. parallel loss of vision in diverse cave fauna (6)). Our study provides the first direct evidence for the hypothesis that flight poses a selective pressure on the speed of olfactory receptor neurons. Our findings also emphasize the energetic costs of rapid olfaction, and the key role of natural selection in shaping dramatic neurobiological shifts.

**Significance Statement:** Flying insects move fast and have therefore evolved exceptionally quick-acting sensory systems. The speed with which such neurobiological shifts can evolve, however, remains unclear. Under the ‘use it or lose it’ hypothesis, loss of flight should lead to degradation of this fast sensory processing ability. We test for evolutionary reductions in olfactory acuity linked to flight loss, through neurobiological comparisons of flightless versus flighted lineages within a wing-polymorphic insect. Our analyses reveal that newly wing-reduced populations have substantially degraded olfactory acuity, with parallel reductions in this sensory ability detected in independently flightless lineages. These findings reveal that flight poses strong selective pressure for rapid olfaction, and highlight the potential of natural selection in rapidly shaping adaptive shifts in animal sensory systems.

## Introduction

The origin of flight posed novel challenges for animals’ sensory systems, including the need for rapid processing of environmental information. Major sensory shifts were needed to compensate for the fact that flying animals move faster and therefore experience more rapid changes in sensory stimuli (vision: (7, 8); olfaction: (2)). This need for rapid sensing seems to be particularly pronounced for olfaction, because both the speed of wind-borne odor plumes and the frequency of odor concentration fluctuations increase with increasing distance from the ground (9, 10). Accordingly, olfactory receptor neurons of flighted insects can respond to odorants extremely rapidly (within 3 ms and with sub-millisecond precision) (11), and can resolve fast odorant fluctuations (above 100 Hz) (3, 12). While it seems obvious that flight must have generated selective pressure for rapid olfactory transduction (the transformation of olfactory stimuli into action potentials), there is still surprisingly little direct evidence with which to assess this hypothesis (13–17). This lack of research on the evolutionary lability of temporal acuity of olfaction stands in stark contrast to the many studies on the evolution of odor specificity, which demonstrate rapid evolutionary adaptation of the olfactory system to the animals’ chemical environments (13, 18–25).

While broadscale evolutionary trends in sensory systems are well documented, the pace at which such systems can evolve is poorly understood. Darwin (26) observed that loss of nonfunctional phenotypes (reductive evolution) is a repeated phenomenon in nature (27, 28). Dramatic examples of such reductive evolutionary processes include rapid deterioration of inactivated genetic material (e.g. (29)), and parallel losses of pigmentation and eyes in diverse cave fauna (e.g. (6, 30)). If maintenance of rapid olfactory transduction is key to the success of flighted insects, this begs the question of what happens to this sensory capacity when such lineages become secondarily wing-reduced. Indeed, flight loss has evolved independently in nearly every order of winged insects, and in some clades it has occurred repeatedly (5, 31).

Under the ‘use it or lose it’ hypothesis, we propose that olfactory acuity becomes rapidly degraded in insect lineages that no longer use flight, in the same way that wings themselves become vestigialized. Here, we use a wing-dimorphic member of the early-diverging winged insect order Plecoptera as a model to test the hypothesis that rapid olfaction is a requirement specifically for flighted insects, but not for flightless lineages. The New Zealand *Zelandoperla fenestrata* Tillyard stonefly complex comprises both full-winged (flighted) and wing-reduced (non-flying) lineages that co-occur widely (32, 33) (Fig. 1A). Recent genomic analyses of this species complex indicate that wing reduction has evolved recently (likely within the last 15000 years) and independently in different regions (4), making this a replicated model system for testing the hypothesis that flightless lineages exhibit secondarily reduced temporal olfactory acuity relative to that of flighted lineages.

**Fig. 1.**
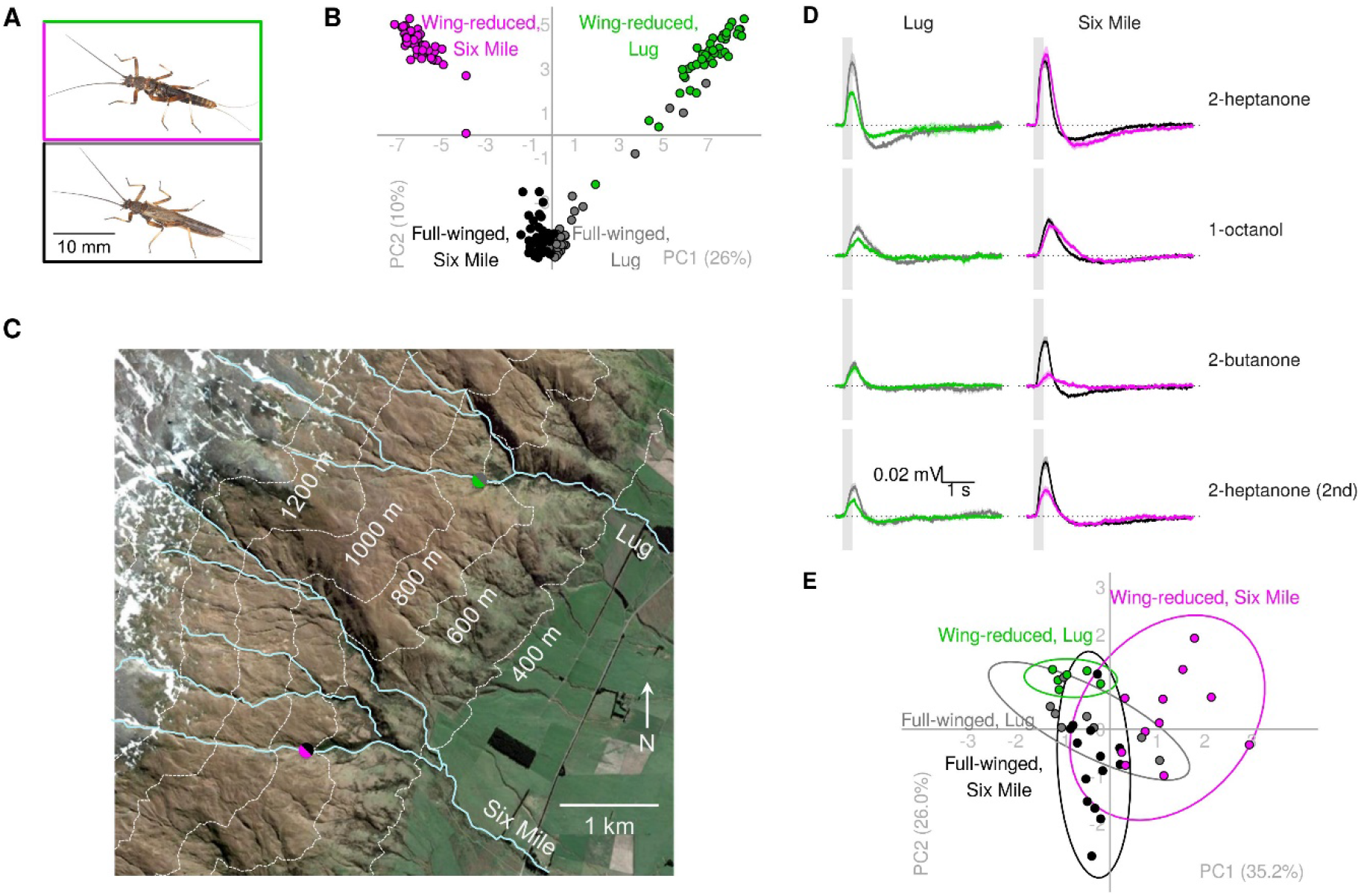
Odorant-evoked antennal responses in full-winged and wing-reduced stoneflies. **(A)** Full-winged (black/grey) and wing-reduced (magenta/green) *Zelandoperla fenestrata* ecotypes. **(B)** Principal component analysis demonstrating the parallel genome-wide divergence between wing-reduced (magenta/green) and full‐ winged (black/grey) ecotypes from Six Mile Creek and Lug Creek (modified from (4)). **(C)** Topographic map of sampling sites (magenta/black and green/grey points) in the Rock and Pillar Range, New Zealand. **(D)** Antennal signal traces of full-winged and wing-reduced stoneflies from Lug (grey: full-winged, 8 antennae; green: wing-reduced, 6 antennae) and Six Mile (black: full-winged, 14 antennae; magenta: wing-reduced, 11 antennae) during stimulation with different odorants (300 ms pulses). Mean ± SEM over antennae. Grey vertical bars indicate valve opening time. **(E)** Principal component analysis demonstrating greater differences in antennal response patterns between wing-reduced (magenta/green) ecotypes from Six Mile Creek/Lug Creek than between full‐winged (black/grey) ecotypes. Ellipses show 95% confidence areas.

## Results and Discussion

To test the hypothesis that flight loss leads to degradation of temporal olfactory acuity, we compared odorant-evoked antennal responses, measured with electroantennogram recordings (EAG), between co-occurring flighted and flightless stonefly lineages (Fig. 1A).

### Odorant-evoked antennal responses are weaker and slower in wing-reduced stoneflies

Antennal recordings in flighted versus flightless stonefly ecotypes from two genetically independent populations (Lug Creek and Six Mile Creek (Fig. 1B, C)) revealed substantial differences in the strength and dynamics of antennal responses to odorants (Fig. 1D, Fig. S1, Fig. S2). The difference in the pattern of responses (strengths, onsets and offsets of responses to 2-heptanone, 1-octanol and 2-butanone) between the Lug and Six Mile populations were greater in the wing-reduced than in the full-winged ecotype (Fig. 1E). These greater differences in response patterns between the wing-reduced stream populations mirror the greater genomic divergence between wing-reduced populations (Fig. 1B). In the Lug Creek sample, wing-reduced individuals showed weaker responses to the odorants 2-heptanone and 1-octanol; whereas in the Six Mile Creek population, wing-reduced individuals showed weaker responses to 2-heptanone and 2-butanone stimuli (Fig. 2A). In addition to weaker antennal responses, wing-reduced individuals from the Six Mile Creek population showed slower response onsets and offsets relative to full-winged individuals (Fig. 2B, C). Neither full-winged nor wing-reduced stonefly antennae could resolve 10-Hz odorant fluctuations (Fig. S1, S2), indicating that antennal responses in stoneflies have a lower temporal resolution than antennal responses in more derived insect species (12, 34, 35), including honey bees (3) (Fig. S4).

**Fig. 2.**
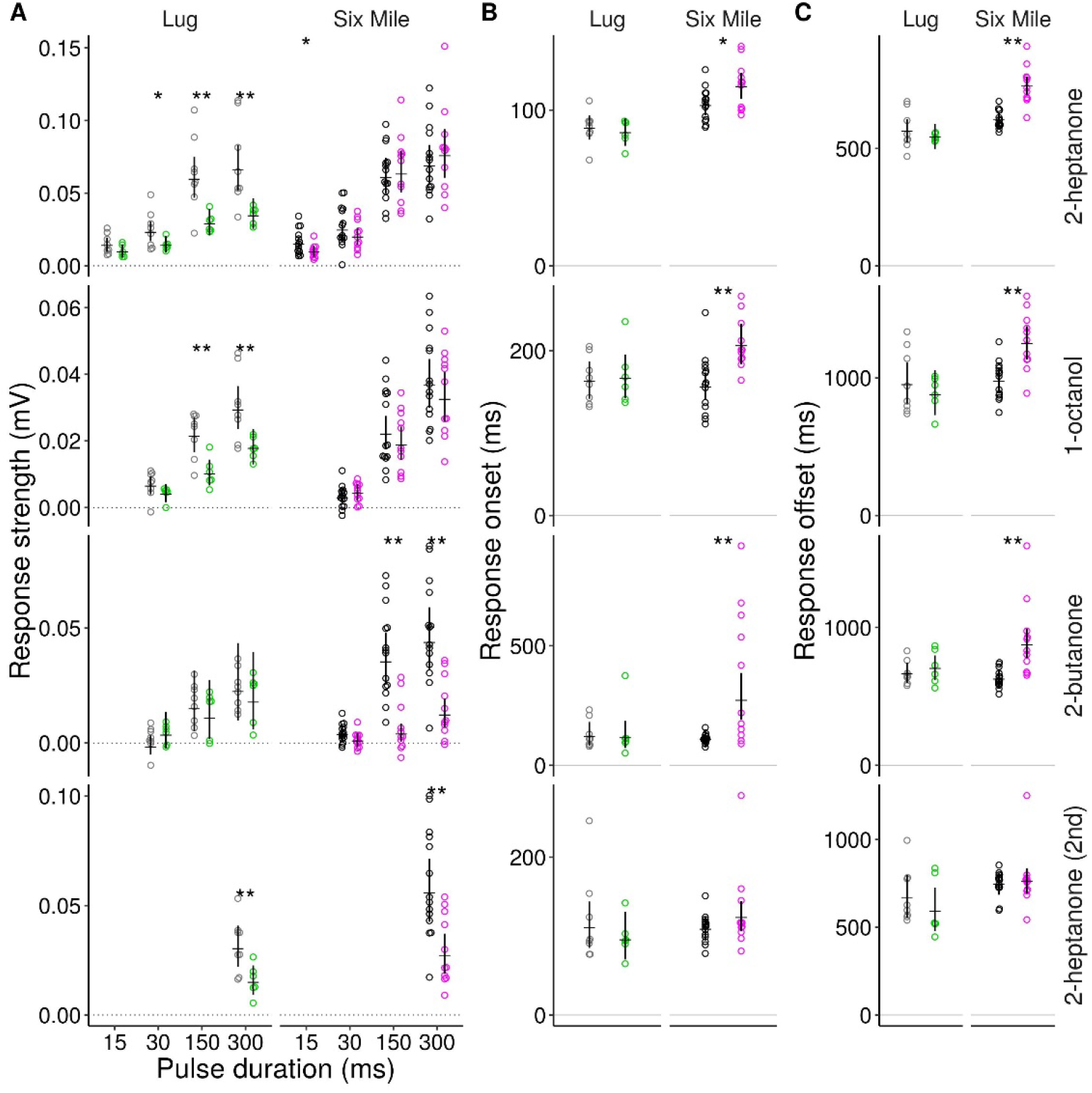
Odorant-evoked antennal responses are weaker and slower in winged-reduced than in full-winged individuals. **(A)** Antennal response strength, **(B)** response onset time (time to 10% of signal maximum after valve opening, includes the time odorant need to reach the antenna), and **(C)** response offset time (time to 10% of signal maximum after the maximum). Black/grey: full-winged; Magenta/green: wing-reduced. Horizontal black lines show means and vertical black lines show 95% credible intervals. *: greater than 95% probability, **: greater than 99% probabilities for differences between antennae of full-winged and wing-reduced stoneflies.

### Weaker antennal responses in wing-reduced stoneflies indicate lower olfactory sensitivity

The weaker antennal responses in wing-reduced individuals suggest these lineages have reduced olfactory sensitivity. Indeed, previous studies have shown that antennal (EAG) responses weaken with decreasing numbers of responding olfactory receptor neurons (36), and with decreasing amplitude of receptor current (37). However, the strength of an antennal response can also be affected by the electrical properties of the antenna itself (37). Therefore, the weaker antennal responses in wing-reduced individuals could reflect a difference in the antennae’s electrical properties.

To test whether weaker antennal responses in wing-reduced stoneflies could reflect non-biological differences between antennae, we recorded antennal responses in live versus dead antennae (Fig. S3A). To induce antennal responses in these live and dead antennae, we applied blank (empty vial), 2-heptanone, 1-octanol, propionic acid, 2-butanone and water (same procedure as in the recordings from live antennae before). Live antennae responded to all odorants. Dead antennae responded to propionic acid and water, but not to the other odorants (see Methods for a discussion of signal source). The strength of odorant-evoked signals in dead versus live antennae were not dependent (Fig. S3B). This lack of dependence confirms that the differences in antennal responses between wing-reduced and full-winged stoneflies reflect true biological differentiation. Therefore, the weaker and slower antennal responses in wing-reduced individuals confirm that flightless lineages have degraded sensitivity and temporal resolution of olfaction.

Given that these sympatric full-winged and wing-reduced ecotypes have similar genetic backgrounds (4, 38), our study provides direct evidence for secondary degradation of olfactory acuity associated with flight loss, while eliminating potentially confounding ecological and evolutionary factors that might also affect olfactory processing.

The finding that wing-reduced stoneflies from Lug and Six Mile Creeks show distinct forms of olfactory degradation (primarily related to response strength in the former, and primarily speed in the latter) (Fig. 2) suggests that these neurobiological shifts evolved independently in each population. This inference is reinforced by genomic analysis (4) confirming independent evolutionary origins for these neighboring flightless populations (i.e. parallel wing-reduction events; Fig. 1B). The current study thus provides additional evidence for their convergent evolution, with different forms of reduced olfactory acuity detected in these independently flightless lineages.

### Selective pressure for rapid olfactory transduction in flying insects

The faster onsets and offsets of odor-evoked antennal responses detected in full-winged stoneflies are likely to facilitate long-range search behaviour. When searching for an odorous target (e.g. a mating partner), flying insects encounter rapidly fluctuating odor plumes (39–41). Because the randomness of air flow destroys most of the directional information present in an odor plume (e.g. concentration gradients), flying animals need to use reactive or infotactic strategies for odor plume tracking, both of which require the animal to rapidly detect and react to the target odor (42–46). A fast onset of the antennal response would shorten the time to detect and react to the target odor, while fast offset of the antennal response would facilitate the detection of subsequent odor stimuli. In addition to facilitating odor plume tracking, rapid olfaction enables the perceptual segregation of mixed odorants from different sources. For example, flying insects can segregate mixed odorants from different sources by detecting millisecond short differences in the arrival of different odorants (47–53).

In line with the ecological requirement for rapid olfaction, insect olfactory receptor neurons can respond to odorants rapidly (within 3 ms) (3, 11). Olfactory receptor neurons can respond rapidly because olfactory receptors are ion channels. These olfactory receptor channels are composed of odorant-specific olfactory receptors (OR) and co-receptors (Orco) (54–56). Missbach et al. (14) suggested that the OR/Orco system represents a specific adaptation associated with insect flight (but see (13) for an opposing view). While OR genes likely evolved in the common ancestor of modern insects, perhaps as an adaptation to terrestrial conditions (13, 25), the OR/Orco system may have arisen more recently (15). Indeed, the fact that genomes of early-diverging (non-flying) insect lineages, such as jumping bristletails (Archaeognatha), have ORs but not Orco, suggests the OR/Orco system has a later origin and facilitates rapid olfaction in winged insects (15).

### Energetic costs of rapid olfactory transduction and mechanisms of its slackening

The reduced strength and speed of antennal responses in wing-reduced stoneflies indicates that olfactory transduction is energetically costly, which in turn generates a selective pressure for slackening olfactory transduction in flightless insects. The strength of an antennal (EAG) response correlates with the receptor current amplitude and action potential number (36, 37). Because, ion currents and action potentials are energetically costly (57), the energetic cost of antennal responses increases with signal strength. Likewise, a rapid antennal response is likely to be energetically costly, because a rapidly responding receptor neuron requires a low membrane time constant so that receptor currents can produce faster rising membrane potentials. The lower membrane time constant, in turn, requires a higher membrane conductance, which increases energy consumption (58, 59). Therefore, the reduced strength and speed of antennal responses in wing-reduced stoneflies could reflect a reduced number of receptors or receptor neurons, a reduced membrane conductance and/or an increased action potential threshold.

Besides fast activation, high temporal resolution requires fast deactivation of olfactory receptor neurons, so that they can respond to following odorant pulses. Deactivation is thought to be mediated by removal of odorants from the OR/Orco complex through odorant-binding proteins and odorant-degrading enzymes (60). Interestingly, a putative odorant-removing protein, *Pinocchio* (61), is significantly more expressed in the notum of wing-reduced than in full-winged stoneflies from Lug Creek (62). If *Pinocchio* is similarly overexpressed in the antennae of wing-reduced stoneflies, this may well explain the weaker odor-evoked antennal responses detected in wing-reduced stoneflies. Future comparisons of antennal expression of olfactory-related proteins, and the number, membrane conductance, and action potential threshold of olfactory receptor neurons will further elucidate the mechanistic basis of this olfactory degeneration.

### Conclusions

Recent analyses have highlighted that shifts in animal mobility can have drastic evolutionary consequences (63). The findings of the current study highlight not only the need for rapid olfactory processing in flying insects (2), but also that this sensory ability can be rapidly degraded when no longer required (i.e. when flight ability is lost). The locally and independently wing-reduced lineages analyzed here are thought to have diverged from their winged counterparts only very recently in evolutionary terms (i.e. during the current interglacial, less than 15000 years ago) (4). This rapid reductive evolution of sensory ability echoes the rapid vestigialization of wings themselves, and also represents a neurobiological parallel to the rapid phenotypic shifts seen under sharp selective gradients in other systems (e.g. loss of vision in cave fauna) (6, 30). Broadly, these findings emphasize the energetic costs of rapid sensory processing ability, and the key role of natural selection in shaping neurobiological shifts. Additionally, this multidisciplinary analysis highlights the potential for future studies to further elucidate dynamic sensory evolutionary processes in the wild.

## Methods

### Stonefly sampling

We sampled stoneflies from zones of ecotypic overlap from two genetically independent (4) stream populations on the Rock and Pillar Range, Otago: Lug Creek (45°25.13’S, 170°7.18’E) and Six Mile Creek (45°26.60’S, 170°5.91’E) (Fig. 1C). Final instar nymphs were collected by hand from under stones or wood in stream cascades and rapids. Nymphs were subsequently reared in the laboratory in Styrofoam cups at 11°C under a natural day:night cycle, in water from their natal stream. After emerging as adults, stoneflies were sexed based on genitalia, and morphologically characterized as either full-winged or wing-reduced.

### Antennal responses to olfactory stimuli

We used electroantennograms (EAGs) to measure the electrophysiological responses of detached antennae to odorants. For the experiments shown in Fig. 1, 2, 3, S1 and S2, we used 1- to 5-day old adult male stoneflies. For the experiments shown in Fig. S3 we used male and female stoneflies. We fixed the stoneflies with adhesive tape and used a razor blade to cut off a 5 mm long section of the distal antennae (the length of the intact antennae was approximately 15 mm). Antennae were mounted with conductive gel (GEL+, Ritex, Germany) on a four-channel silver electrode (64) (Fig. 3A). To eliminate between-session variability (e.g., due to humidity or circadian rhythms in antennal responsiveness (64)), the left and right antennae of one full-winged and one wing-reduced stonefly were recorded simultaneously (Fig. 3B). Five minutes later, the antennae were placed at a distance of 2 mm in front of the outlet of the olfactory stimulator.

**Fig. 3.**
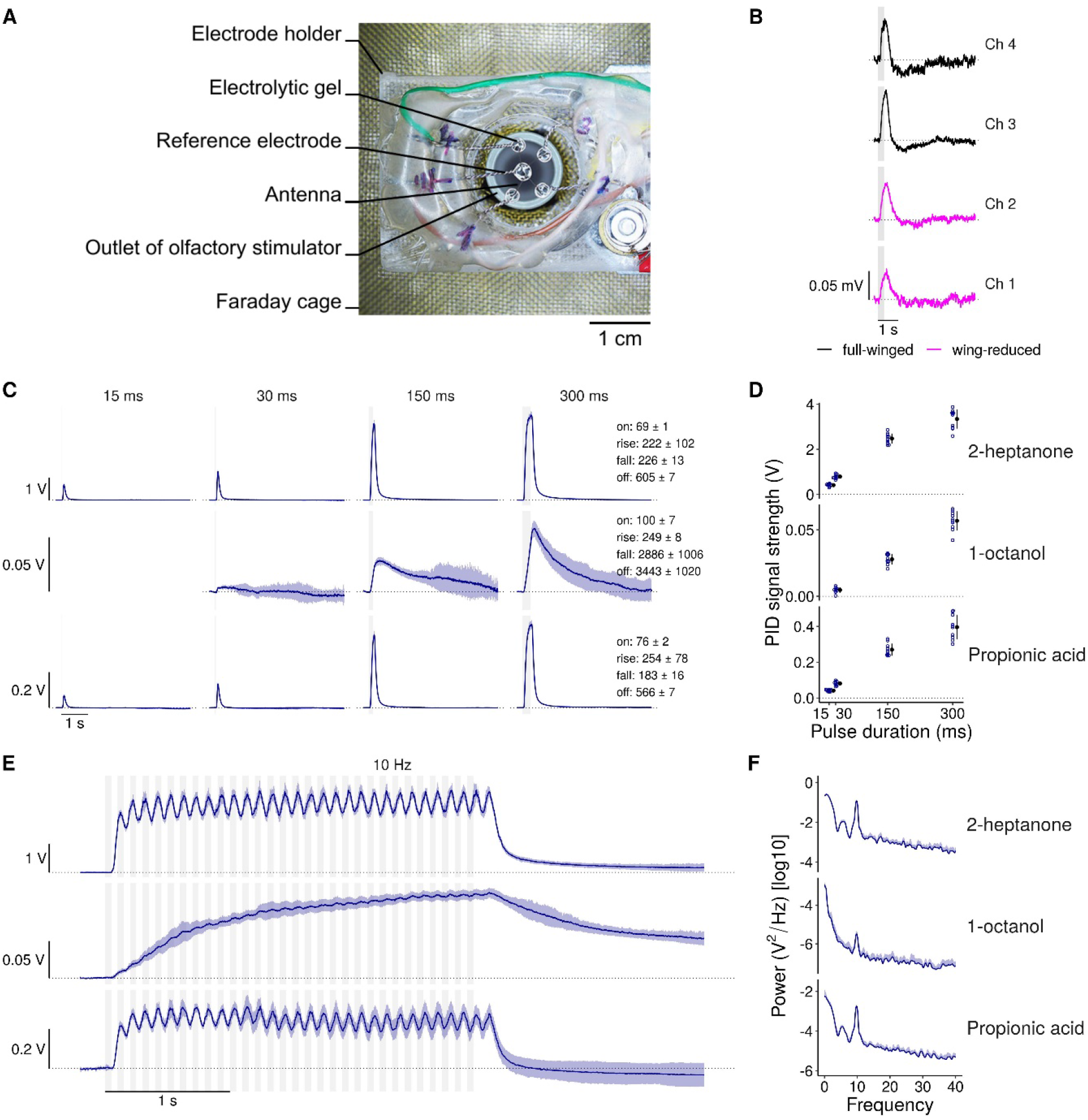
Experimental setup, example EAG signals and olfactory stimulus dynamics. **(A)** Experimental setup with the quadruple electrode and outlet of the olfactory stimulator in a Faraday cage. **(B)** Example EAG signals of four stonefly (Six Mile) antennae simultaneously recorded (median over 10 stimuli of 300 ms long 2-heptanone pulses). **(C)** Photoionisation detector (PID) measurements of odorant stimuli (mean ± SD across 10 recordings). We did not measure 2-butanone, because this odorant saturated the PID. Values show response parameters of 300 ms-pulses: Onset time (on, time to reach 10% of the signal maximum after valve opening, includes the time odorant need to reach the sensor), rise time (rise, 10% to 90%), fall time (fall, 90% to 10%), and offset time (off, time from maximum valve opening to 10% after signal maximum). All times are given in ms (mean ± SD). **(D)** PID signals strength (mean ± SD). **(E)** Same as C) but with fluctuating 10-Hz stimuli. **(F)** Power spectral densities of the EAG signals during 10-Hz stimulation (mean + SD).

EAG signals were differentially amplified against the reference electrode using 1000x gain, recorded in AC-coupled mode and low-pass-filtered at 1 kHz (MA 103 preamplifier and MA 102 four-channel amplifier, Universität zu Köln). The distal tips of the four antennae were mounted on a common central electrode that was connected to the inverted inputs of the preamplifiers. Under this approach, positive EAG signals reflect excitatory responses (activation of olfactory receptor neurons) and negative EAG signals reflect inhibitory responses.

### Olfactory stimulation

We used a custom-made 6-channel olfactory stimulator that was designed to minimize mechanical stimulus components (same approach as in (65) but built with materials as described in (66)). This stimulator provided a constant airflow (volume flow rate = 4.8 L/min, flow speed = 100 cm/s). The antennae were exposed to this constant airflow throughout the whole duration of the recordings before and between the odorant stimuli. To apply odorants, odorant-laden air (300 mL/min) was injected into the constant carrier air stream and, simultaneously, the same amount of clean air was withdrawn to keep the total airflow constant. The odorant air stream was produced by bottled compressed air and the carrier air stream was produced by an aquarium pump. Both air streams were filtered with active carbon filters (HN4S-AUN, Parker). The stimulation was controlled with the data acquisition system Micro 3 1401 and Spike2 software version 8.03 (CED).

All odorants were kept in glass vials with PTFE septum screw caps (20ml EPA vial, JG Finneran). We presented the following olfactory stimuli in the following sequence: odorless blank (empty vial), 2-heptanone, 1-octanol, propionic acid and 2-butanone (these four odorants purchased from Sigma-Aldrich), and female stonefly. We chose the odorants for the following reasons: 2-heptanone because it induces robust antennal responses in different insect species (3); 1-octanol because it was used in a previous EAG study of stoneflies (67), 2-butanone because of its fast stimulus dynamics (48), and the headspace air above a virgin female stonefly, to test for antennal responses to a possible female sex pheromone (note that we recorded antennal responses of unmated males that have not encountered female pheromones before). We also trialed propionic acid, which had evoked particularly strong antennal responses in a previous study (67), but we ultimately excluded this odorant following the detection of non-biological artifacts (Fig. S3; see below).

We used a photoionization detector (miniPID B, Aurora) to measure the dynamics of odorant concentration change (Fig. 3 C–F). To vary the strength of olfactory stimuli we presented each odorant at up to four different pulse durations (15, 30, 150 and 300 ms valve open time). Because the rise time of the odorant concentration is larger than 150 ms (Fig. 3C), these different stimulus durations result in different maximum concentrations. Note that odorant-specific differences in stimulus dynamics are a consequence of interactions between odorants and the surfaces of both the olfactory stimulator and the photoionization detector (66, 68).

We also recorded antennal responses in honey bees (Fig. S4) as a scale for comparing stoneflies’ antennal responses to other insects (see temporal resolution of antennal responses of honey bees, locusts, moths, and cockroaches in a previous study (3)).

Each olfactory stimulus (odorant/pulse duration combination) was presented 10 times at an inter-stimulus interval of 5 seconds. Ten seconds after the last 300 ms long stimulus we presented fluctuating 10-Hz stimuli by repetitively opening the valve for 50 ms at a frequency of 10 Hz over a period of three seconds (Fig. 3E–F). This 10-Hz stimulus served to quantify the temporal resolution. Five seconds after the 10-Hz stimulus the next odorant started. At the end of each experiment we presented another 10 stimuli of 2-heptanone (300 ms) to test whether the antennae are still responding. Each recording session lasted approximately 16 minutes.

### Testing the effect of non-biological electrical antenna properties on antennal responses

To identify any potential non-biological EAG signals, we recorded 9 additional stonefly antennae (from 5 male and 4 female full-winged individuals from Six Mile). After the recording, we killed the antennae by applying 90°C hot water vapor for 2 to 5 min prior to another set of EAG recordings with the same stimulus protocol. To test whether the response to stonefly odor (Fig. S1) could be due to humidity we presented water.

While the propionic acid and water induced negative EAG signals both in live and dead stonefly antennae, other odorants showed no signals in dead antennae (Fig. S3A). Therefore, while EAG signals evoked by propionic acid and stonefly odor may include non-biological components, the other odorants assessed (2-heptanone, 1-octanol, 2-butanone) have no such artifacts. Propionic acid-evoked responses in dead antennae could be explained by dissociation of propionic acid when it comes in contact with water vapor in the air. This creates H^+^ and OH^−^ ions, and uneven accumulation of these ions at the recording and reference electrodes would induce electrical potentials.

To test whether the signal strength of live antennae is dependent on their physical properties, we utilized the negative signals in dead antennae induced by propionic acid (physical property) and compared them to the response strength to 2-heptanone (because this odorant induced the strongest responses) of the same antennae, but when still alive (Fig. S3B). A linear relationship would indicate that the difference in response strength between full-winged and wing-reduced stonefly antennae could result from differences between the physical properties of antennae themselves. To test for such differences, we performed linear regression analyses for each of the different pulse durations (15, 30, 150, and 300 ms). We used the response strength to 2-heptanone in nine live antennae as dependent variable and corresponding response strength to propionic acid of the same nine antennae when dead as independent variable. We assessed whether the slope in each of the four regression analyses where significantly different from zero. The finding that the strength of odorant evoked signals in live antennae did not depend on the signal strength evoked in dead antennae (Fig. S3B) supports the biological rather than physical source of response differentiation between full-winged and wing-reduced individuals.

### Data Analysis

EAG signal data were exported from Spike2 and then further analyzed using R (version 3.6.3) (69). Each recording was cut into 5 s traces, starting 0.2 s before and ending 4.8 s after each stimulus onset. Each trace was baseline-corrected by subtracting the median voltage during the 0.2 s time window before valve opening from the entire trace. To increase the signal-to-noise ratio, we calculated the median trace over each set of 10 stimulations with the same odorant/pulse duration combination. To reduce the noise, we applied a running median filter with a window size of 11 ms on the traces.

To assess whether an antenna shows stimulus-induced responses to the last set of 2-heptanone stimulations, we defined a response threshold as two times the standard deviation of the last 3 s of the trace. If the stimulus induced response of an antenna to these 2-heptanone stimulations did not exceed the response threshold for at least 50 consecutive ms between 100 and 800 ms after valve opening, we excluded all recordings from this antenna from further analysis (22 of 24 full-winged antennae and 17 of 24 wing-reduced antennae were further analyzed).

We quantified different response parameters evoked by 2-heptanone, 1-octanol, and 2-butanone between stonefly ecotypes for a given odorant and collection site: 1) response strength as the mean response in the time window of 25 ms before and 25 ms after the maximum signal between 50 ms and 1s after odor valve opening, 2) response onset: time between valve opening and 10% of maximum signal (before signal maximum), 3) response rise time: time between 10% to 90% of maximum signal (before signal maximum), 4) response fall time: time between 90% to 10% of maximum signal (after signal maximum), 5) response offset time: time between valve opening and 10% of maximum signal (after signal maximum), and 6) temporal resolution for 10-Hz stimulations: power spectral densities on a 3 s time window starting 0.4 s after the first valve opening using the *multitaper* R package (70) with the *sine taper* method.

To quantify if there are differences in response strength between ecotypes, we ran linear mixed models with log2 transformed response strength as the response variable. To avoid negative values, we added an offset of 0.01 to each mean response prior to the log2 transformation. We included ecotype (full-winged or wing-reduced) and pulse duration as explanatory variables, and added an interaction between both variables. To account for repeated measurements of the same antennae, we included antenna identity as a random factor.

For data of the second set of 300 ms long 2-heptanone stimuli, we used a linear model with the response as the response variable and ecotype as the only explanatory variable.

To test if there are differences in response timing, i.e. in response onset time, rise time, fall time or offset time between stonefly ecotypes for a given odorant and collection site, we ran linear models. We included the log2 transformed response timing of interest (either response onset, rise time, fall time, or response offset) for 300 ms long pulses as the response variable and ecotype (full-winged or wing-reduced) as explanatory variable.

Inferences for all types of models were drawn using Bayesian statistics. We used an improper prior distribution (flat prior) and simulated 10.000 random draws from the posterior distribution using the function *sim* from the R package *arm* (71).

We used the 50% quantile as the mean and the 2.5% and 97.5% quantiles as the lower and upper limit of the 95% credible interval. We calculated the proportion of simulated values from the posterior distribution that are bigger for one ecotype (e.g. A) over the other (e.g. B). A resulting proportion of 0.99 would mean that we are 99% certain that ecotype A has a larger parameter value than ecotype B. For plotting, we back-transformed the quantiles in the original scale. For the mean response strength, we subtracted the offset of 0.01 again. We marked differences between full-winged and wing-reduced ecotypes in a given odorant/pulse duration configuration of >95% certainty with 1 asterisk and >99% certainty with 2 asterisks.

To visualize the differences in odorant response patterns across full-winged and wing-reduced antennae of stoneflies from Lug and Six Mile Creeks, we performed a principal component analysis (PCA). We selected the response parameters response strength, onset time and offset time for stimulations with 300 ms long pulses of 2-heptanone, 1-octanol, 2-butanone and 2-heptanone (2nd). We used each of the odorant / response parameter combinations as a single input variable for the PCA (4 odorants × 3 response parameters = 12 input variables).

## ACKNOWLEDGEMENTS

The photoionization detector was funded by a University of Otago Research Grant (#3435, #18817) to PS, and J.M.W. and G.A.M. were partly funded by a Marsden contract (UOO1412). We thank Michael Thoma for comments on the manuscript and for pointing us to the possibility that propionic acid evokes non-biological EAG responses, and we thank the participants of the Behavioral Ecology and Evolution Seminar (Dept. Zoology, University of Otago) for feedback on the manuscript.

## Author contributions

Conceptualization, P.S. and J.M.W.; Methodology, P.S. and S.N.; Software, S.N.; Validation, S.N. and P.S.; Formal Analysis, S.N.; Investigation, P.S.; Resources, S.N., G.A.M., B.J.F., J.M.W., and P.S.; Data Curation, S.N. and P.S.; Writing – Original Draft, P.S, J.M.W.; Writing – Review & Editing, P.S., S.N., J.M.W., G.A.M., and B.J.F.; Supervision, P.S.; Funding Acquisition, P.S.

---

**Fig. S1.**
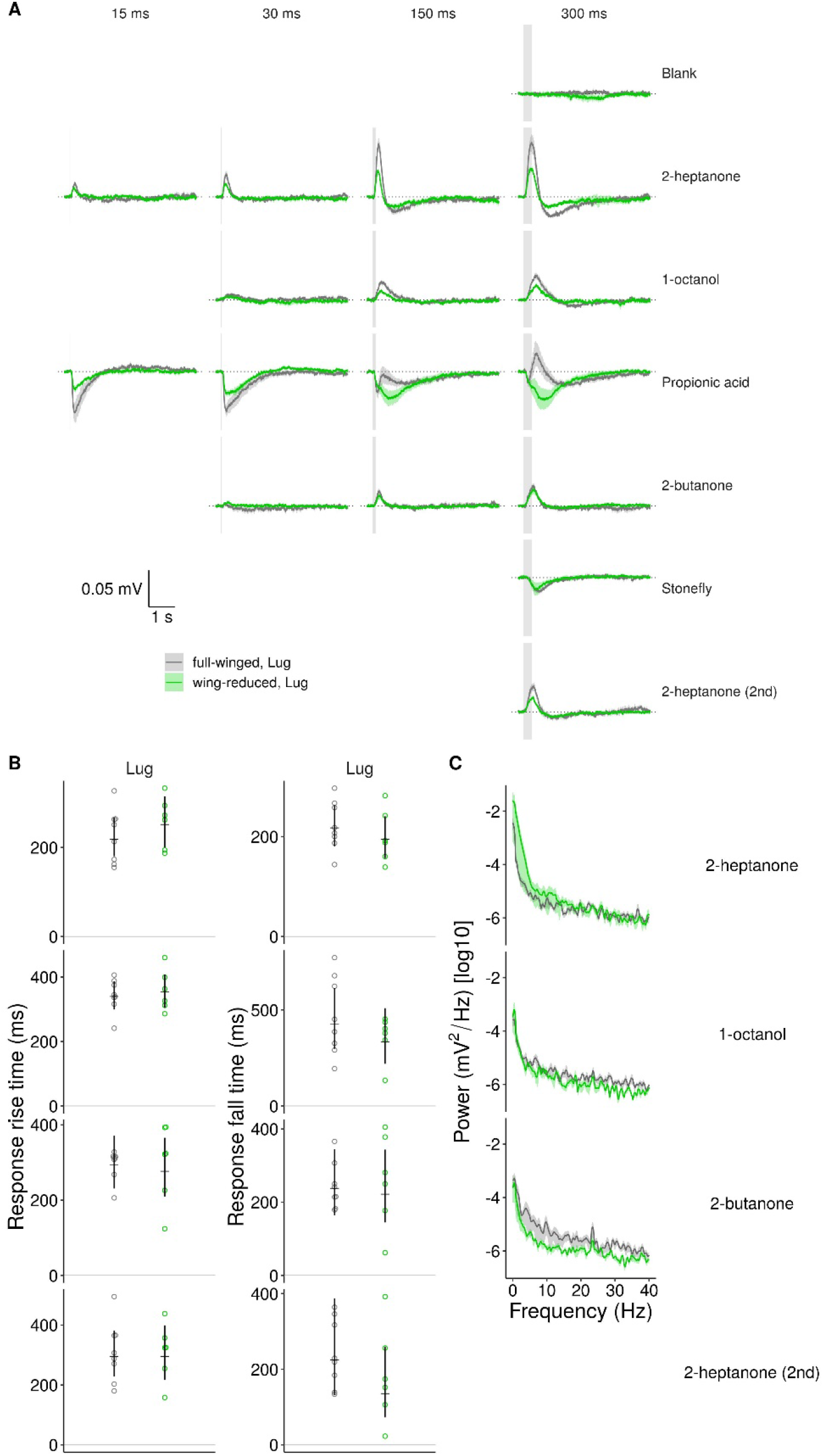
Dynamic properties of antennal signals (Lug Creek). **(A)** Antennal signal traces of full-winged (grey) and wing-reduced (green) individual antennae during stimulation with different odorants (rows) and pulse durations (columns). Mean ± SEM over antennae. Grey vertical bars indicate valve opening time. The sequence of panels (top left to bottom right) correspond to the sequence of stimuli. Note that signals evoked by propionic acid and stonefly may contain non-biological artifacts (see Fig. S3A). **(B)** Rise time (time between 10 and 90% of signal maximum) and fall time (time between 90 and 10% of signal maximum) for Lug Creek population. **(C)** Power spectral densities of antennal signals in response to fluctuating 10-Hz stimuli (Lug Creek population). The lack of peaks at 10 Hz indicates that the antennal responses of stoneflies could not resolve odorant fluctuations at a rate of 10 Hz.

**Fig. S2.**
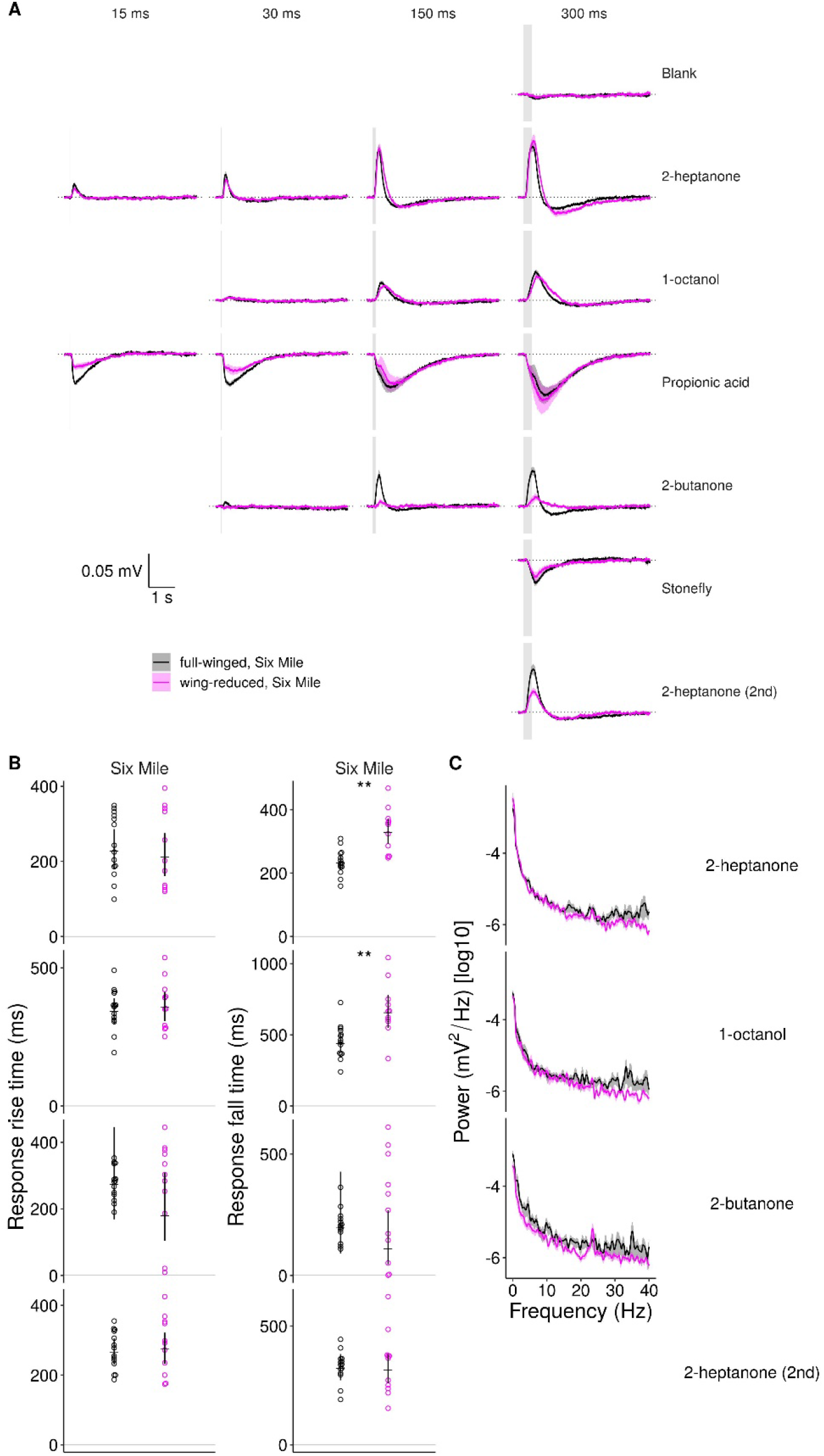
Dynamic properties of antennal signals (Six Mile Creek). Same as in Fig. S1 but for Six Mile. Black: full-winged; magenta: wing-reduced.

**Fig. S3.**
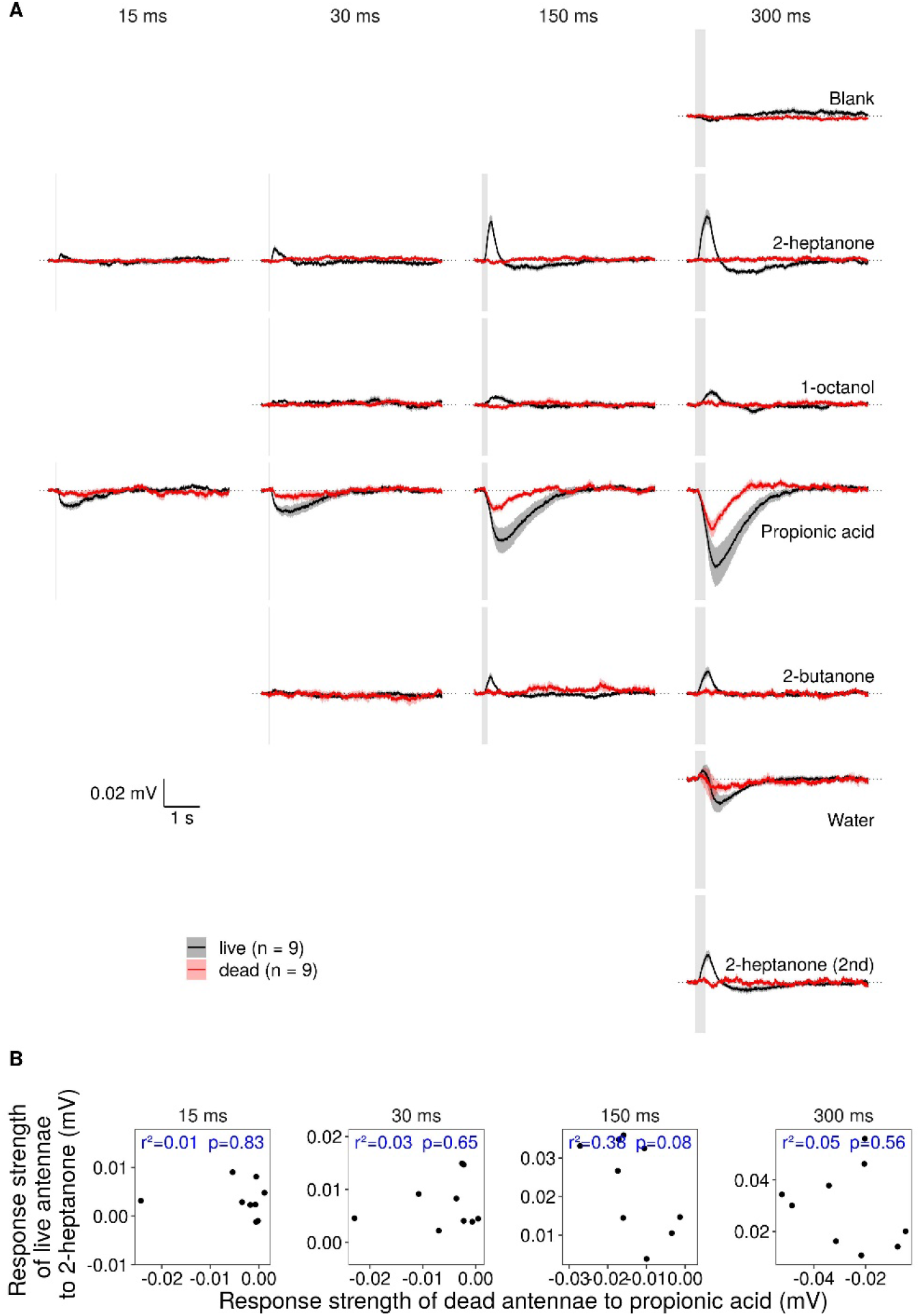
Biological and non-biological antennal signals are independent from each other. **(A)** EAG signal traces of live (black) and dead (red) antennae (full-winged stoneflies, dead and alive antennae were identical) during stimulation with different odorants (rows) and pulse durations (columns). Mean ± SEM over antennae. Grey vertical bars indicate valve opening time. The sequence of panels (top left to bottom right) corresponds to the sequence of stimuli. **(B)** Scatter plot of signal strength in dead antennae to propionic acid in relation to their signal strength to 2-heptanone when these antennae were still alive. Each dot per panel shows an individual antenna. Blue values show the proportion of variance (r^2^) explained and the p-value (p) for non-zero association between dependent and independent variable.

**Fig. S4.**
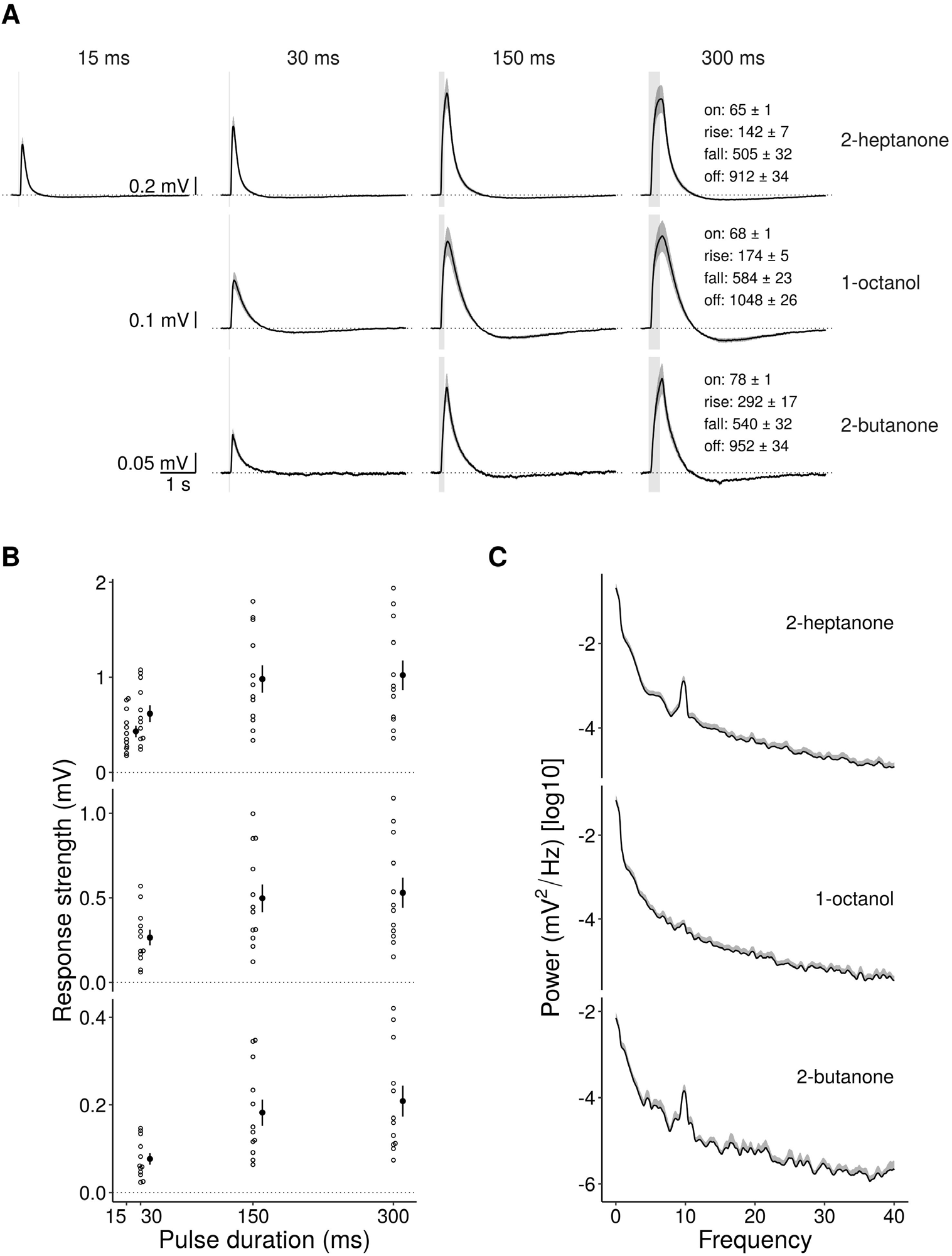
Odorant-evoked responses of honey bee antennae. **(A)** EAG signal traces from honey bee antennae during stimulation with different odorants (rows) and pulse durations (columns). Mean ± SEM over antennae. Grey vertical bar indicates valve opening times. Values represent: Onset time (on, time to reach 10% of the signal maximum after valve opening, includes the time an odorant needs to reach the sensor), rise time (rise, 10% to 90%), fall time (fall, 90% to 10%), and offset time (off, time from maximum signal to 10% of maximum). All times are given in ms (mean ± SEM). Data from 12 antennae. **(B)** EAG signal strengths (mean ± SEM). **(C)** Power spectral densities of the EAG signals during fluctuating 10-Hz stimuli (mean + SEM).

